# Advancing protein display on bacterial spores through an extensive survey of coat components

**DOI:** 10.1101/2024.11.22.624950

**Authors:** Todd C. Chappell, Trevor B. Nicks, Jan-Fang Cheng, Angela Tarver, Jessica A. Lee, Arushi Kalia, Nikhil U. Nair

## Abstract

The profound stability of bacterial spores makes them a promising platform for biotechnological applications like biocatalysis, bioremediation, drug delivery, etc. However, though the *Bacillus subtilis* spore is composed of >40 proteins, only ∼12 have been explored as fusion carriers for protein display. Here, we assessed the suitability of 33 spore proteins (SPs) as enzyme display carriers by direct allele tagging at native genomic loci. Of the 33 SP investigated, 26 formed functional fusions with β-glucuronidase (GUS) – a ∼272 kDa homotetramer. This almost triples the number of SPs assessed for enzyme display and doubles the number of functional fusions documented in the literature. We quantitatively assessed the 1) SP promoter activation dynamics during growth, 2) supported GUS activity on spores, 3) surface availability, 4) protection from thermal and proteolytic degradation, and 5) compatibility of co-expression of many of these fusions. Multi-copy expression and pairwise co-expression of the most promising SP-GUS fusions highlighted the complexity of spore structure/assembly and difficulty in predicting compatibility between different SPs fusions. We also assessed the suitability of engineered spores to degrade PET (polyethylene terephthalate) films and found that surface exposed SPs were most effective. Beyond the broad survey, a key outcome of our work was the identification of SscA (<u>s</u>mall <u>s</u>pore <u>c</u>oat assembly protein <u>A</u>) as an effective spore display carrier. SscA supported enzyme activity at least fourfold higher than any other SP, including the well-established and popular anchor, CotY. We attribute this to the biochemical and genetic features of SscA; its promoter demonstrated early and sustained activation relative to other SPs and its small size (∼3 KDa) likely minimally interferes with enzyme folding, oligomerization, and activity. Although the specific localization of SscA within the spore coat has not been established, its hydrophobic nature and low surface availability suggests that it assembles internally within the spore coat, which makes it highly stabilizing and suitable for many biocatalytic applications. Overall, this work serves as a knowledgebase to advance the biotechnological utility of *B. subtilis* spores.

## INTRODUCTION

Surface display technologies are highly versatile, genetically-encoded protein presentation platforms used to characterize, engineer, and deliver proteins. In addition to simplifying manipulation and substrate/target accessibility, surface fusion can enhance the stability of the displayed protein against a variety of denaturants, making surface display attractive for a range of biotechnological applications including biocatalysis, drug delivery, bioremediation, etc. In particular, bacterial (endo)spore display has garnered interest for the deployment of proteins in more challenging or extreme conditions due to the robust nature of spores. Spores remain intact and viable under intense thermal, pH, protease, and solvent stressors, and some of this stability can be passed on to displayed proteins.

To enable spore display, the protein of interest is genetically fused to a component of the proteinaceous shell that surrounds the spore core^1–3^. One of the most studied sporulating is the model Gram-positive bacterium *Bacillus subtilis*. Detailed localization and interaction studies of proteins expressed during sporulation have revealed the *B. subtilis* spore to be a complex structure of more than 40 proteins organized into morphologically distinct layers including the: i) basement layer, ii) inner coat, iii) outer coat, and iv) crust. While many spore proteins (SPs) have been successfully fused to fluorophores to determine protein localization, only a limited number have been investigated for functional applications of spore display. Most SPs investigated for display applications are crust (CgeA, CotV, CotW, CotX, CotY, CotZ) or outer coat proteins (CotA, CotB, CotC, CotD, CotG), while a limited number of studies have also explored the use of the inner coat protein OxdD^4–6^. Though these studies hint at the robustness of spores and the versatility of different components to support display, most SPs remain unexplored for their suitability for spore display.

In this study, we sought to provide a broad, functional characterization of SPs as fusion partners in order to provide guidance on the design of spore display systems. After a thorough analysis of potential SP fusion partners, we developed a plate-based screening method for allelic tagging of SPs with an enzyme reporter at their native loci via natural co-transformation/congression. We characterized the most active fusions ability to i) assemble onto spores, ii) support enzymatic activity, iii) be surface accessible, iv) stabilize fusion partners against thermal and proteolytic inactivation, iv) express in multicopy and co-express together, and v) sporulate efficiently. Our initial results highlighted a previously understudied protein, SscA, as a highly promising candidate for spore display. By exploring the activity of combinations of the most active fusions, we also show that optimal combinations of spore display fusions cannot be determined by simply selecting the most active individual fusions. Finally, we explored the accessibility of our top fusions partners to larger substrates by displaying the plastic-degrading enzyme DuraPETase on the spore surface and demonstrating functional degradation of PET films. Cumulatively, this work describes the most extensive and direct comparison of different SPs for their suitability to spore display and will aid in the design of better spore display constructs for a variety of biotechnological applications.

## RESULTS

### A plate-based screening method facilitates identification of scarless allelic fusions of spore proteins (SPs)

Most *B. subtilis* spore display studies integrate fusion constructs at neutral genomic loci (e.g., *amyE, lacA, thrC*, etc.), which result in two copies of the SP – the fusion and the wildtype (WT)^3^. However, spores assembled from such designs may not accurately reflect SP fusion potential due to competition between copies of SPs and expression from non-native genomic contexts – both of which can alter spore coat composition and/or integrity. To mitigate these issues, we decided to assess the suitability of each SP (the carrier) by directly tagging it at its native genetic locus with a reporter (the cargo). After identifying more than 50 known spore coat proteins, we narrowed the candidate field by removing any SPs previously identified as critical for proper spore assembly or known to impede sporulation as a fusion (**Table S1**). For the remaining 44 SP candidates identified (**Figure S1**), we designed a theoretical spore display library of 88 N- and C-terminal integration cassettes that consist of 2 kb homology arms flanking the β-glucuronidase gene (*uidA*, GUS), a GSSSG spacer linking uidA to the SP, and a 6x-His tag on the free terminus of *uidA* (**Supplementary Data File 1**). To ensure scarless tagging, we leveraged congression during natural transformation in *B. subtilis*. After co-transforming the fusion constructs along with an unlinked selectable marker (linearized pBS1K/C) that integrates at an unlinked locus (*amyE*), we directly plated the cells on sporulation agar containing X-gluc (5-bromo-4-chloro-3-indolyl-β-D-glucuronic acid) for blue/white screening (**Figure 1A, B, Figure S2**).

**Figure 1.**
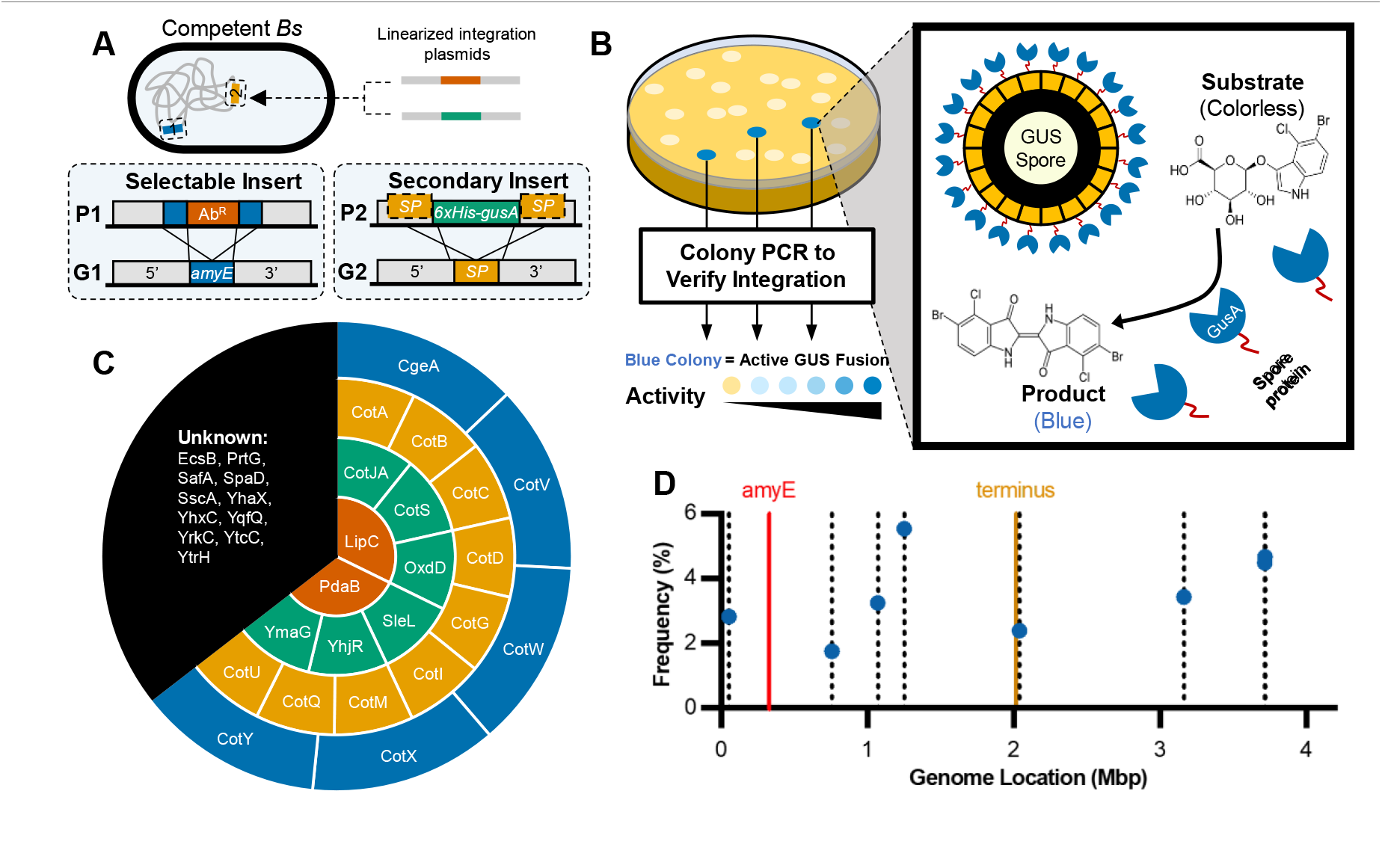
Construction of GUS tagged spore protein (SP) library using co-transformation. **A)** Schematic of co-transformation process and integration vector design. Linearized plasmids P1 carries a selectable antibiotic cassette with flanking homology to target integration at the *amyE* locus. Linearized plasmid P2 encodes N- or C-terminal fusions of *gusA* to SP gene with 2 kb homology arms for scarless allelic tagging. **B)** Screen for GUS^+^ colonies where blue colonies in the plate screen indicate functional expression of SP-GUS fusion. **C)** Location of the 33 unique spore proteins that could be confirmed by blue-white screening and PCR; crust (blue), outer coat (yellow), inner coat (green), cortex/basement layer (orange) and unknown location (black). **D)** Co-transformation frequencies ranged between 1.5 – 6% for the 8 loci used to assess factors affecting congression. The location of *amyE* and *ter* are also indicated for reference. The location of all 44 loci under investigation are in **Figure S1**.

Of the 88 possible constructs designed, we were able to successfully synthesize 52 integration vectors, which represented at least one N- or C-terminal fusion for 33 of the original 44 SPs (**Figure 1C**). On screening these 52 vectors for GUS activity on sporulation plates, we found 34 constructs (∼65%) produced blues colonies and could be verified as integrated at the expected locus by PCR. This corresponds to 26 unique spore proteins, of which 9 have uncharacterized localization and the remaining are distributed across the 4 distinct layers of the spore: i) cortex/basement layer, ii) inner coat, iii) outer coat, and iv) crust. C-terminal fusions accounted for 23 of the 34 active GUS fusions, and 16 fusions were to SPs that have not been previously used for spore-display of enzymes (**Table S2**).

Collectively, co-transformation efficiency (efficiency = blue colonies/Ab^R^ colonies) ranged considerably for the 34 integration vectors. We further evaluated 8 of these constructs (targeting loci spread throughout the *B. subtilis* genome) to identify any features that may contribute to the differences in the observed co-transformation efficiencies (**Figure 1D**). Although congression has previously been reported to correlate with the distance from the selected marker and replication fork, we did not find any such trend in our studies. We also found no correlation with the size of the spore protein being targeted or the gene orientation. Analysis of the sense DNA strand suggested that the co-transformation efficiency was modestly correlated positively to the A (adenosine) content and negatively to the G (guanosine) content (**Figure S3A–B**). Further, it was strongly positively correlated to the A+C content (**Figure S3C**). Since altering A+C content of the sense strand does not affect annealing interactions with its complementary strand, we investigated the Gibbs free energy of the most stable predicted secondary structure formed by the integrations cassette and found that it correlated negatively with co-transformation efficiency (**Figure S3D**). Since DNA enters the cytoplasm in single-stranded form, our data suggests that highly stable secondary structure formation may adversely affect integration efficiency.

### Allelic tags provide insight into SP promoter dynamics

To determine relative expression and activation timing of the different SP promoters, we performed time-course studies of GUS activity on sporulation plates for all 34 fusion constructs that produced blue colonies (**Figure 2A–F**). Though the observed activity of N- and C-terminal fusions of PdaB – the only SP from the cortex/basement layer – were relatively weak, strong GUS expression could be detected from protein within all other layers of the spore. SPs could be generally divided into three main categories: (I) very low activity, which are either poorly expressed and/or form inactive fusions with GUS; (II) late expression/low activity, which exhibit activity after an extended incubation; (III) early expression/high activity, which show significant activity early that is retained during extended incubation (**Figure 2G**). Half (17) of the tested constructs fall within (III). Interestingly, the location within the spore did not correlate with any of these three categorizations, which suggests that promoter strength and tolerance to GUS fusion are stronger determinants of activity than localization within the spore.

**Figure 2.**
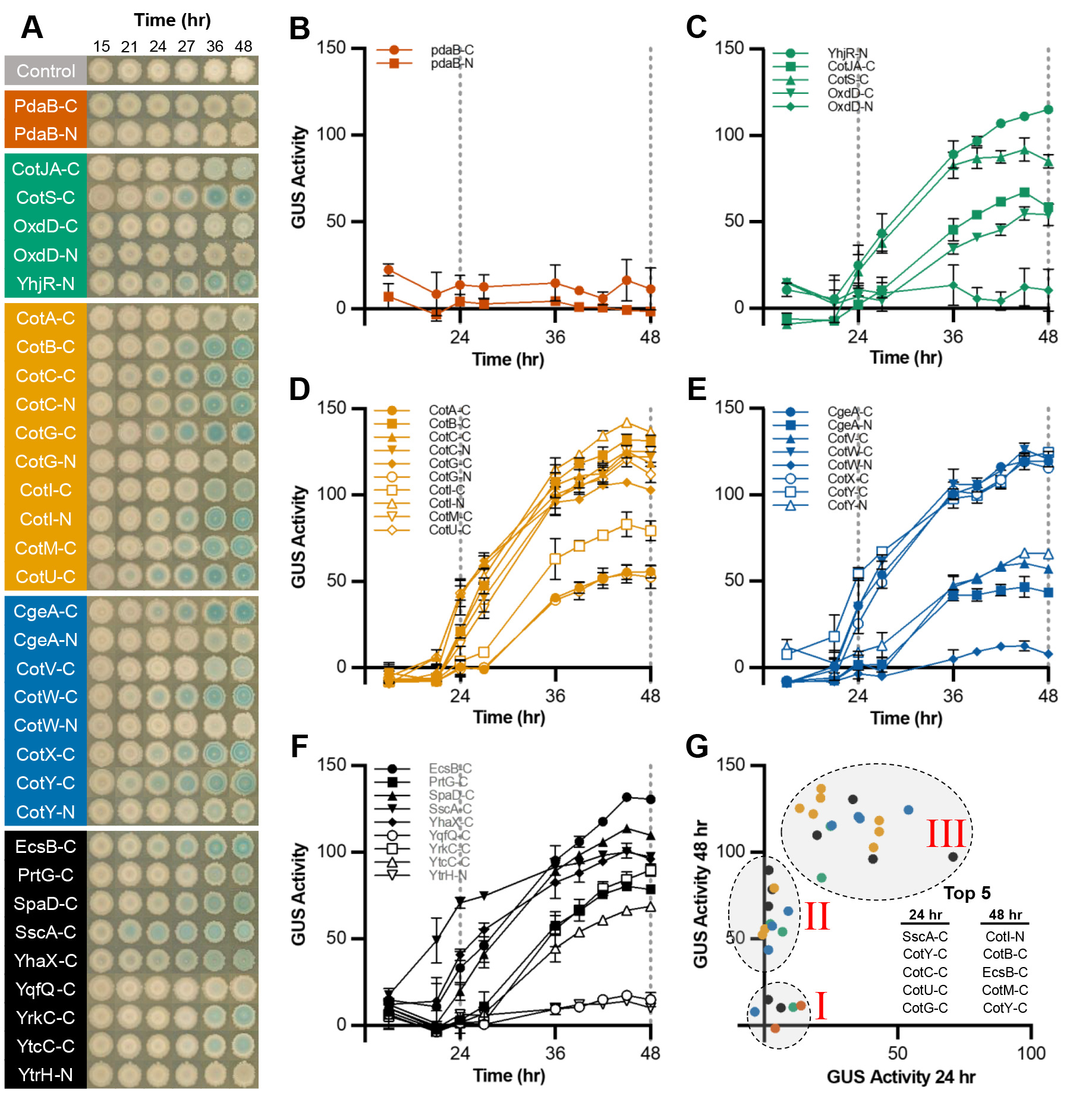
Activity dynamics of SP-GUS fusions under screen conditions. **A)** Time-course images of colonies of the 34 *B. subtilis* strains with confirmed GUS-tagged SP alleles (WT – grey; cortex/basement – orange; inner coat – green; outer coat – yellow; crust – blue; unknown location – black). Quantified time-course GUS activity determined by image analysis of microcolonies in (A) for **B)** cortex/basement, **C)** inner coat, **D)** outer coat, **E)** unknown location SP-GUS fusions. Data points are the average of 2 biological replicates and error bars are SEM. **G)** Scatter plot of 24 h and 48 h average GUS activity. SP-GUS fusions can demonstrate (I) low activity, (II) activity during late sporulation, (III) activity during early and late sporulation. The five most active fusions are listed in inset “Top 5” for both 24 and 48 h timepoints.

### Characterization of GUS activity on spores identifies SscA as a promising carrier for biocatalysis

While the plate assay effectively determined relative promoter activation and expression of active SP fusions, it did not indicate whether the fusions are properly incorporated into spores. Detectable GUS activity can be attributed to enzyme derived from vegetative cells, actively differentiating cells, and/or lysed mother cells that were not incorporated into spores. To assess spore-specific GUS activity, we generated spores in liquid cultures and thoroughly washed them to remove non-specifically bound proteins. We found that basement/cortex fusions were inactive on spores as well, whereas all other fusions yielded some degree of activity (**Figure 3A**). The highest activity fusion partners with known locations are in the outer coat and crust of the spore. Specifically, CotY, CotG, CotX, and CgeA were 4 of the 5 best performing fusion partners, which supports their general popularity as fusion partners in the published literature {REF}. Interestingly, SscA – a small protein of ∼3 kDa with unknown location that has not previously been assessed as a carrier for spore display – yielded the highest activity among all SPs tested, with more than 4-fold higher activity relative to the next most active fusion partner, CotY.

**Figure 3.**
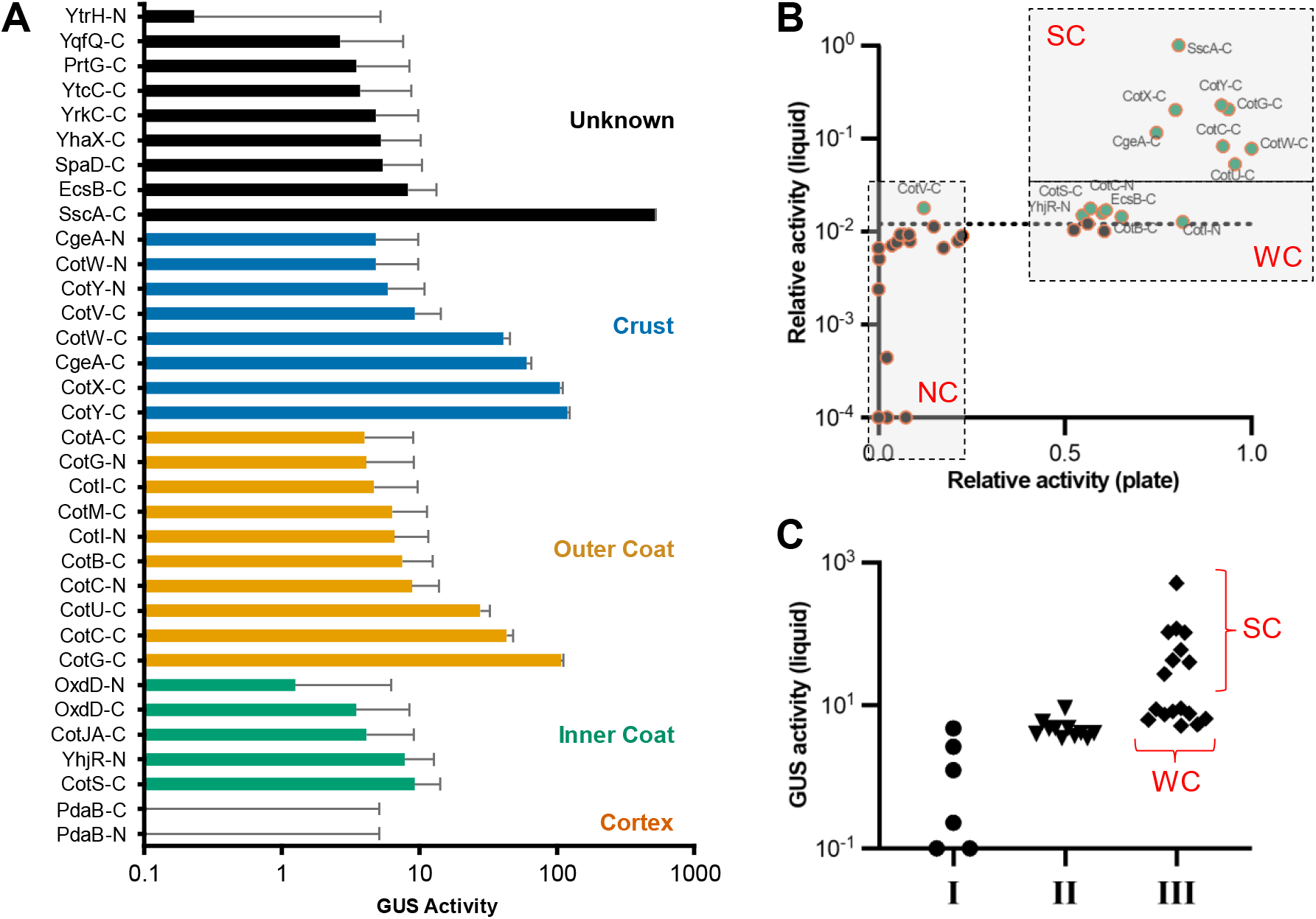
Relative activity of spore displayed SP-GUS fusions. **A)** Activity of SP-GUS fusions that have assembled onto spores. Data represents an average of 6 biological replicates and error bars are SEM. **B)** Correlational plot of relative GUS activity on solid and liquid media. Activity on agar plates is the slope of the average activity of fusions from 15-27 h indicated in Fig. 2B-G, and that from liquid culture is the end datapoint shown in (D). NC are negative candidates, WC are weak candidates, and SC are strong candidates. **C)** Plot of liquid culture activity per the activity categories defined in Fig. 2G. Both WC and SC exhibit activity designated as category (III) on plate assays.

To determine the effectiveness of the plate assay to identify candidates for spore display, we compared the GUS activity on plates (determined using slope during early growth, 15-27 h) and that of isolated spores (derived from liquid cultures). Both data sets were normalized to the maximum, and values below the limit of detection were set to 0.0001 to accommodate the log scale. From this analysis, fusions could be categorized into 3 groups: i) strong candidates (SC), which have high activity in, both, plate and washed spores preparations, ii) weak candidates (WC), which demonstrate high activity on plates but low activity as isolated spores, and iii) negative candidates (NC), which have low activity as, both, colonies on plates and as washed spores (**Figure 3B**). We did not find any candidates that demonstrated low activity on plates but high activity as spores, highlighting the robustness of the plate-based screen. We found that all SC and WC are category (III) of the plate screen (**Figure 3C**), which represent 50 % (17/34) of the fusion candidates, 47 % (8/17) of which are SC. This demonstrates that the plate-based screen is a reliable approach to eliminate inactive or low activity fusions proteins.

### Crust proteins fusions are most surface accessible

Surface accessibility is an important metric for certain spore display applications, such as antigen presentation and polymer degradation. To assess surface accessibility using immunostaining, we wanted to isolate highly pure spore. However, when we attempted to purify spores from cultures using the standard lysozyme treatment approach^7^, we found it produced insoluble aggregates of cell debris that bound the secondary antibody and distorted results (**Figure S4**). So, we developed an alternative spore purification method that relies on the separation of spores and cells via differential centrifugation. Our method makes use of the differences in spore and vegetative cell size and densities. Centrifugation generates a layer of loosely packed spores on top of a tightly packed bed of cells and forespores (**Figure 4A inset**). Using purified spores, we characterized surface accessibility of the SP-GUS spores by labeling them with two different primary antibodies that target the 6xHis tag or GUS, respectively, in independent experiments (**Figure 4A**). In general, the polyclonal anti-GUS antibody provided higher surface detection levels than the monoclonal anti-6xHis antibody, likely due to its ability to probe multiple epitopes in a complex spore assembly where any single epitope could be occluded due to packing/steric hindrances. Our results indicate that CotY-C was by far the most surface accessible of the fusions, while SscA-C was largely undetectable. In general, surface availability broadly correlated with our expectation based on published spore structure, with crust and outer coat proteins yielding, on average, stronger signals and basement/cortex proteins (**Figure 4B**). However, there were several notable exceptions. For example, CotW and CotG, which have been proposed to be comprise the outermost layers of the spore^8^ had the lowest signals. Conversely, the inner coat protein OxdD was highly detectable. These results are consistent with observations Potot et al. who noted that an OxdD fusion was more surface exposed than CotG^6^. This suggests that that OxdD may not only be confined to the inner coat. Proteins with previously uncharacterized locations generally exhibited moderate to low surface availability. Interestingly, EcsB-C, PrtG-C, YqfQ-C, and YtrH-C exhibited surface detection equivalent to crust and outer coat proteins, suggesting that they may also assemble at the periphery of spores.

**Figure 4.**
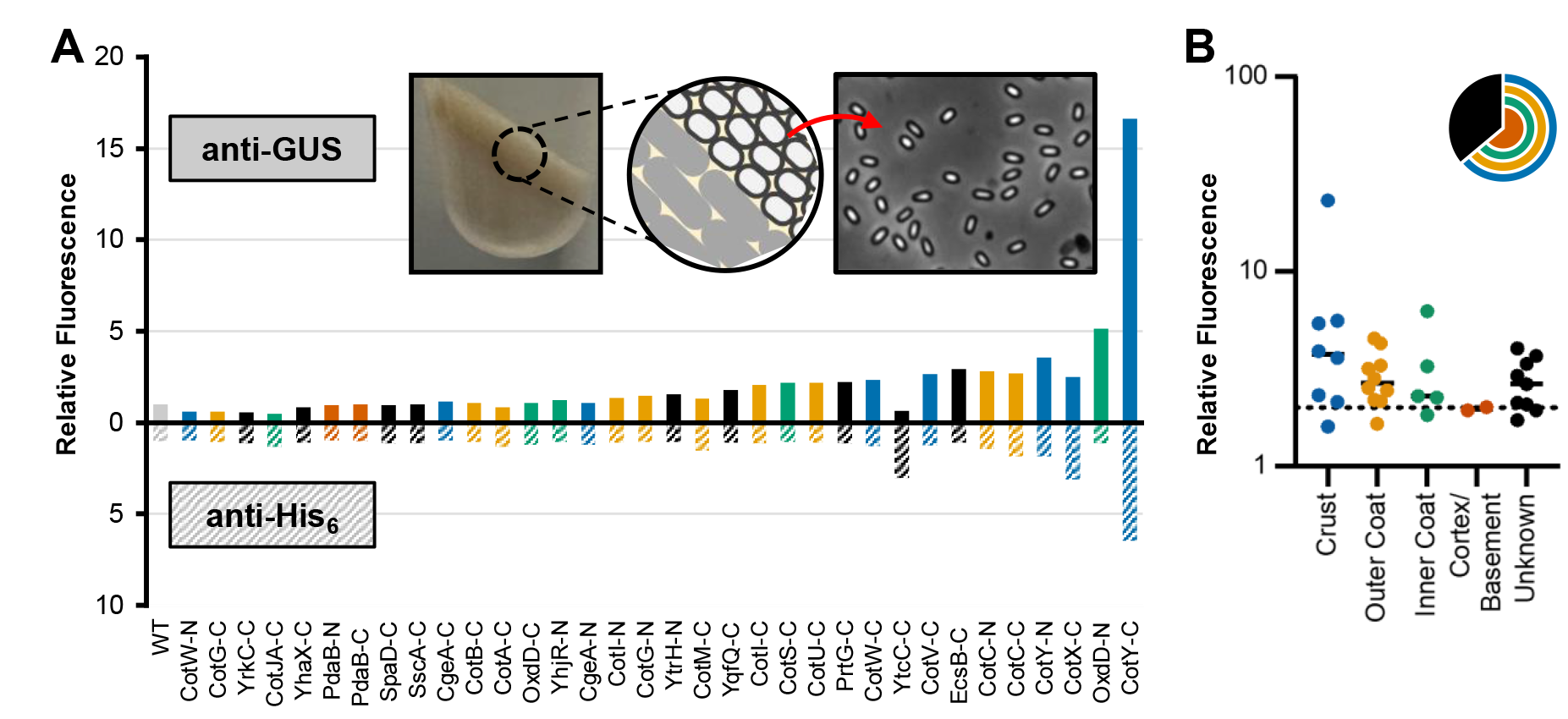
Surface accessibility of SP-GUS fusions. **A)** Mean fluorescence of purified, immunostained spores normalized to WT. Immunostaining was performed using two primary antibodies that target different antigens: anti-6xHis (bottom) and anti-GUS (top). In both cases, a fluorescent secondary antibody was used for detection and spores were analyzed via flow cytometry. Samples are arranged in order from lowest to highest combined fluorescence. Inset shows layering of spores on top of cells in sporulation cultures and validation of spore purity in top layer using phase microscopy at 100× magnification. **B)** Relative fluorescence of samples shown in (A) organized by location in spore. Bars indicate median value for spore layer and show a trend of decreasing detectability when GUS is fused to spore proteins located in deeper spore layers.

### Several SP fusions stabilize GUS against inactivating agents but do not significantly affect sporulation efficiency

Thermal denaturation and proteolytic degradation are common challenges encountered by enzymes during biocatalysis. We investigated how resilient our 15 most active spore fusions were against these challenges. For the thermostability challenge, we incubated purified GUS and SP-GUS spores for 15 min at 37, 50, and 60 °C. While the free enzyme control showed no significant loss of activity at 37 °C, incubation at 50 and 60 °C resulted in only 27 % residual activity and complete loss of activity, respectively (**Figure 5A, Figure S5**). In contrast, all the SP-GUS fusions retained at least 78 % activity after the 50 °C incubation, with most (CgeA-C, CotY-C, CotX-C, CotW-C, CotC-C, CotG-C, CotU-C, EcsB-C, SscA-C) exhibiting elevated activity – a phenomenon commonly observed due to stabilization by surface display^9–12^. Even after the 60 °C challenge, 6 fusions exhibited greater than 50% residual activity. CgeA-C retained the most relative activity (85%) at 60 °C, though SscA still retained more than fivefold higher total GUS activity compared to CgeA-C. Upon challenge with 0.1 mg mL^−1^ trypsin, we also saw elevated activity for most SP-GUS fusions (**Figure 5B**). Increasing trypsin concentrations to 1.0 mg mL^−1^ generally resulted in a loss of activity, though most fusions still retained at least 50 % activity relative to control untreated spores. Further, SscA retained the highest activity at all trypsin concentration, retaining 91 % relative activity even when treated with 1.0 mg mL^−1^ trypsin.

**Figure 5.**
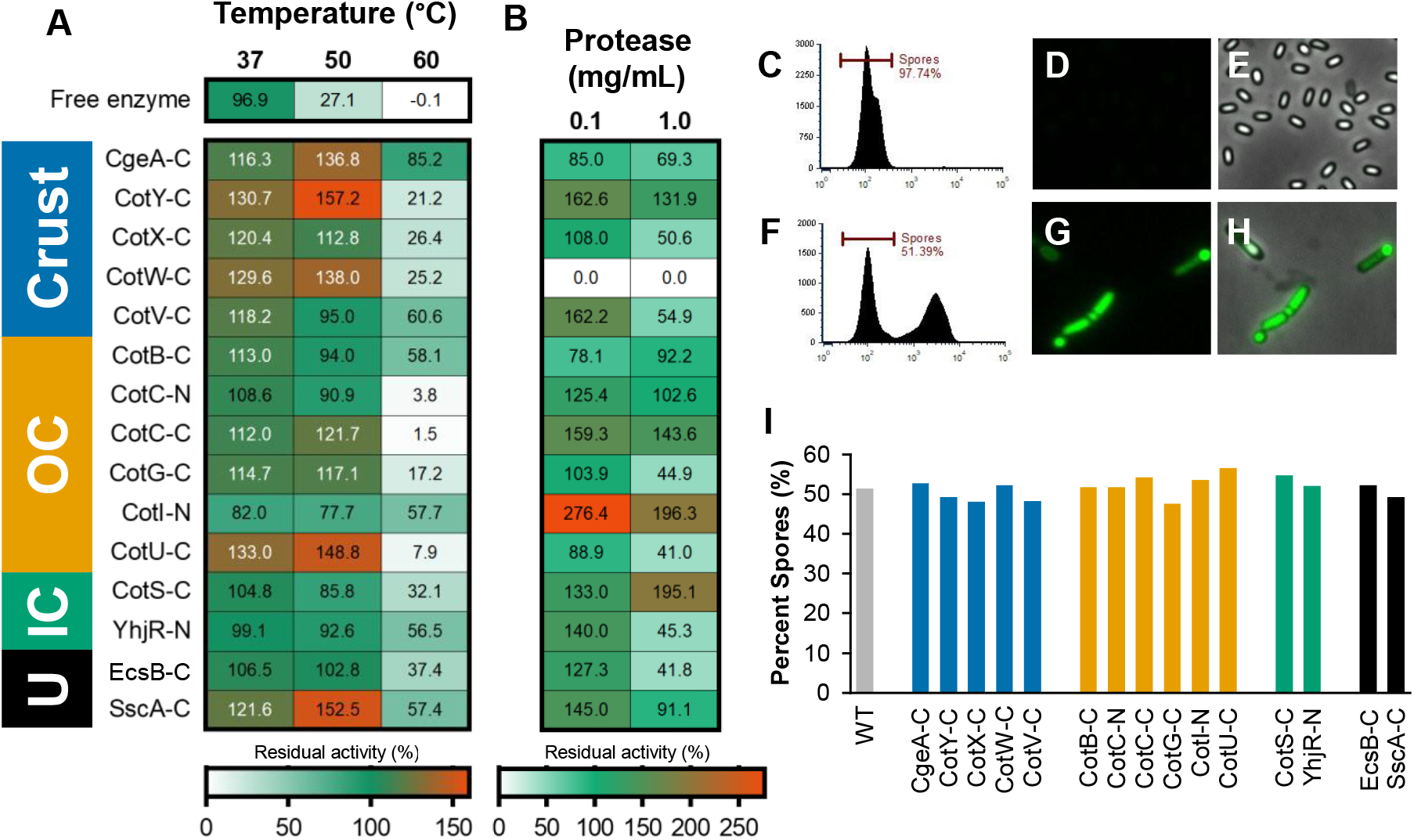
Assessing GUS stability against inactivating challenges. **A)** Residual activity of SP-GUS fusions after 15 min incubation at indicated temperatures The free enzyme rapidly inactivates at elevated temperatures whereas most fusion – especially those on the crust – demonstrate modest thermal activation at 37 and 50 °C, and inactivation at 60 °C. **B)** The free enzyme rapidly inactivates on protease treatment whereas most spore coat displayed fusions are protected. Fusions to crust proteins is least protective, as expected, given their solvent accessibility. For both (A) & (B) activity is relative to initial activity at 37 °C for each construct. **C)** Flow cytometric, **D)** fluorescence, and **E)** phase contrast images demonstrating isolation of purified spores. Cytosolically expressed YFP is localized only to mother and vegetative cells and do not label spores. Conversely, **F)** flow cytometric, **G)** fluorescence, and **H)** phase contrast images of spores and cells shows labeling of non-spore cells. **I)** Sporulation efficiency of all constructs was approximately even. U = unknown location, IC = inner crust, OC = outer crust.

To assess whether modification of spores at their native locus alters sporulation efficiency, we developed a method of differential staining using the DNA-binding dye SYBRSafe. The difference in permeability of the spore and cell membrane permits staining of vegetative and mother cells, while leaving spores unstained (**Figure 5C–H**). Spores purified using our previously described purification protocol showed no fluorescence and phase bright spores (**Figure 5C–E**). Unpurified samples revealed the selective staining of remaining phase dark cells following sporulation (**Figure 5F–H**). Having confirmed that spore and cell fractions can be efficiently distinguished, we assessed sporulation efficiency of SP-GUS fusions using flow cytometry. In all cases. sporulation efficiency ranged from ∼45-55 % indicating only minor changes in sporulation efficiency for SP-GUS expressing strains (**Figure 5I**).

### Co-expression of select SP-GUS fusions highlights complexity of their assembly on spores

Co-expression of multiple fusions could increase display density of GUS and provide insights into compatibility of different SPs for engineering applications. We first investigated the locus dependence and co-loading potential by comparing the activity of the five most active SP-GUS fusions (SscA-C, CotX-C, CotY-C, CotG-C and CgeA-C) when integrated at: i) *amyE* only (*amyEΔ::sp-gusA sp*^*+*^), ii) *amyE* only with the native SP copy knocked out (*amyEΔ::sp-gusA spΔ::erm*), iii) the native locus only (*amyE*^*+*^ *sp::sp-gusA*), and iv) both the *amyE* and native loci (*amyEΔ::sp-gusA sp::sp-gusA*) (**Figure 6A**). All constructs expressed genes using their native promoters.

**Figure 6.**
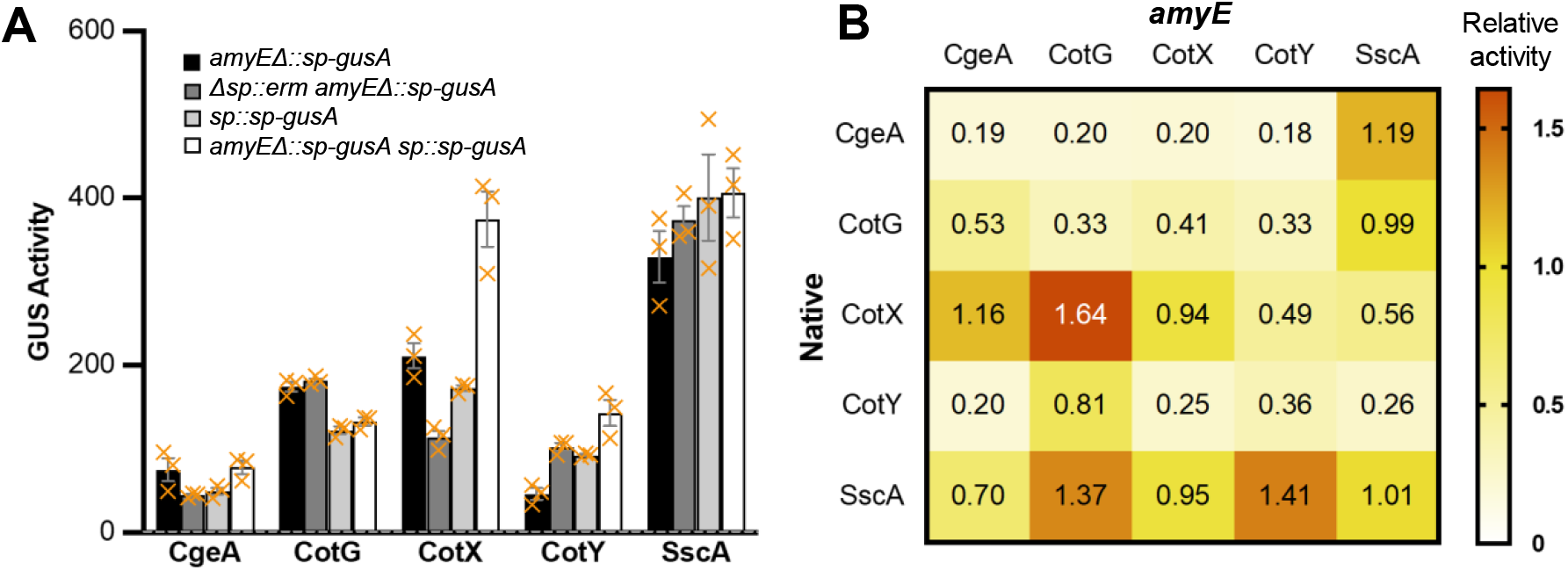
Co-expression analysis of promising spore display candidates. **A)** Assessing the genetic context dependence of SP-GUS fusion activity. The *sp-gusA* fusion gene was expressed after integration at *amyE* locus only, the native locus of the SP, or both. In the case of the first, the native copy of the SP gene was left intact or knocked out. Only fusions to CotX-C and CotY-C showed higher activity when expressed from two different loci. SscA-C fusions demonstrated high activity in all configurations. The locus dependence on the presence of a second WT or tagged SP gene copy was gene-dependent. **B)** Co-expression of two GUS fusions – one from the native locus (*sp*) and the other from *amyE*. SscA-C and CotX-C fusions (at their native loci) generally yielded higher activity when co-expressed with fusions of other proteins expressed from the *amyE* locus. Activities are represented relative to the SscA-C fusion at its native locus.

Interestingly, we found that the fusions exhibited quite varied responses depending on the integration locus and presence of a second copy of the SP gene. The CgeA-C fusion was slightly more active when integrated at the native locus and was not affected by an additional copy of unmodified or *gusA*-tagged *cgeA*. Conversely, the activity of the CotG-C fusion was higher when integrated at the *amyE* locus relative to its native locus. It was not affected by the presence of unmodified *cotG*, and, surprisingly, the activity of the double integrant was lower when compared to a single integrant. The activity of CotX-C was improved in the presence of WT *cotX*, did not differ measurably based on integration locus, and improved markedly when integrated at two loci. The activity of the CotY-C fusion was also higher when present in two copies. However, unlike that of CotX-C, its activity was higher from the native locus relative to that from *amyE*. Interestingly, activity from the *amyE* locus improved when the WT copy was knocked out. Compared to other spore carriers, SscA-C demonstrated the least variation based on locus and copy number and retained high activity with all constructs tested.

We next tested co-expression of two of the top five GUS fusions – one at the native SP locus and the other at *amyE* (**Figure 6B**). In general, fusions expressed from the *amyE* locus often yielded higher activity when co-expressed with CotX-C and SscA-C fusions from their native loci. Specifically, CotX-C and CotG-C were the most synergistic as were SscA-C with CotG-C and CotY-C. Overall, the behavior and interactions of the fusions were very SP-dependent, further highlighting the complexity of spore assembly, structure, and genetic context of associated gene expression.

### Spore-displayed PETase is most active when surface accessible

Although SscA is a promising display anchor for GUS – an enzyme with a small, freely diffusible substrate, we wanted to assess how it and other SPs would support activity of an enzyme with a large insoluble substrate. We selected the PET (polyethylene terephthalate) depolymerizing enzyme, DuraPETase^13^, and generated fusions with the five most promising GUS fusion carrier and integrated them at *amyE*. When we first estimated PETase activity using the small molecule substrate analog *p-*nitrophenol acetate (pNPA), we found that the SscA-C fusion displayed the highest activity (**Figure 7A, Figure S6**). However, when we tested enzymatic activity of the same five engineered spores on PET films, we found that the CotY-C fusion was most active (**Figure 7B**). This is consistent with the high surface accessibility relative to other SPs. The CgeA-C fusion demonstrated relatively low activity, consistent with prior reports by Bartels et al. who found fusion of laccases to this protein to not be very active^5^. Interestingly, SscA still demonstrated moderately high activity, equivalent to, or higher than, other crust proteins like CgeA and CotX – indicating that it may still present enzymes to the surface and a highly versatile anchor for spore display.

**Figure 7.**
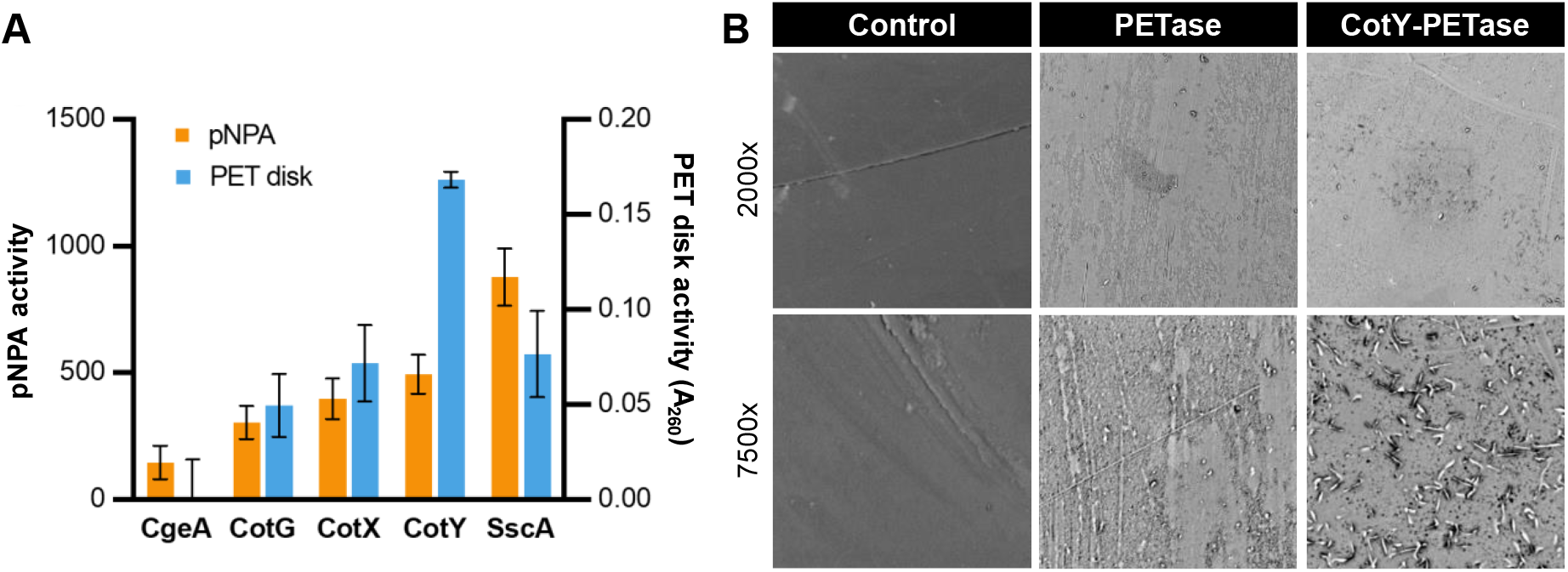
Assessing the activity of SP-PETase fusions using pNPA and PET disks. C-terminal fusions of DuraPETase to CgeA, CotG, CotX, CotY, and SscA were active on **A)** the soluble substrate pNPA (*p*-nitrophenol acetate) and an end-use PET disk. The SscA-C fusion was most active of the soluble substrate whereas the CotY fusion was most activity on insoluble PET. **B)** SEM images of PET films treated with control WT *B. subtilis* spores (left), purified DuraPETase (middle), and *B. subtilis* spores expressing CotY-DuraPETase fusion (right). The unmodified spores we unable to etch the disks, whereas the purified enzyme and recombinant spores visibly modified the PET surface.

## DISCUSSION

The profound stability of bacterial endospores and unique ability to combine protein expression with surface immobilization independent from secretion or soluble-protein purification, positions spores as a powerful tool for diverse applications. In this work, we almost triple the number of SPs tested as carriers for display to 33 and identify 26 functional fusions, of which 16 had not previously been used as display carriers. Although our goal was to assess 44 coat proteins, 9 protein fusions could not be characterized due to technical difficulties with DNA synthesis and strain construction.. We selected GUS as the reporter to evaluate SPs as potential fusion partners for two major reasons – i) ease of characterization and ii) because as a >270 kDa tetrameric enzyme, it is more representative of fusion proteins used in biocatalysis than model monomeric fluorescent proteins and enzymes. One of our key findings was the discovery of SscA as a spore protein (SP) carrier, which is poorly characterized and not documented for engineering applications. The SscA fusion was approximately four-fold more active than other well-characterized best-in-class SPs like CotY and CotG. This could partly be attributed to the small size of SscA, being a 28 aa long peptide, which may minimally interfere with the folding and activity of enzymes like GUS. The early and sustained activation of the SscA promoter may also be a contributor to the high level of activity seen for its GUS fusion. A prior transcriptomic analysis of spores reported that SscA was the only spore coat protein in the top 46 of most abundant mRNA, which also supports our assessment that it is highly expressed^14^. Additionally, CotB, CotG, and CotH were previously found to be significantly reduced in spores derived from a *sscA*Δ background, suggesting SscA may play a key role in spore coat assembly^15^.

In addition to significantly expanding the number of spore display carriers, we used allelic tagging to fuse SPs to GUS rather than choosing to create merodiploid strains. We constructed our strains by leveraging congression during natural transformation whereby unlinked cassettes are concurrently integrated and different sites in the chromosome. This also allowed us to briefly assess factors affecting congression. We found that the melting temperature of the strongest secondary structure within the sense strand of the 2kb long homology arms was inversely proportional to congression frequency. Although this work does not provide any specific proof for this assertion, our work provides a hypothesis that could be tested more rigorously in future studies. In any case, allelic tagging has previously not been used for spore display applications. Based on our results, we found that direct modification of SPs at their native loci can be beneficial in certain cases, like for SscA and CotY, but in other cases, expression from *amyE* was preferred (e.g., CgeA, CotG, CotX). Allelic tagging also enabled us to assess the promoter activation dynamics of all SPs. In general, the best carriers were characterized by early-activating promoters with prolonged expression. Late-activating promoters – identified as those with low GUS activity within the first 24 h – generally yielded spores with relatively low GUS activity. While we cannot altogether preclude that GUS fusions may alter protein stability, folding, etc. and therefore, our assessment of promoter dynamics, our results are nevertheless consistent with the limited literature on comparative strengths of SP promoters and the resulting loading density of spore-displayed cargo proteins^5,16^.

An analysis of all fusion proteins using immunostaining and flow cytometry using anti-6xHis and anti-GUS antibodies revealed that CotY was the most surface exposed protein of those tested. Overall, the signals were higher with the polyclonal anti-GUS antibody. This may be because it is able to bind to several epitopes and would therefore by less sensitive to steric hindrances. Many known crust and outer coat proteins like CotC, CotV, and CotX also demonstrated high surface availability whereas the basement/cortex protein PdaB and others did not. But not all crust and outer coat proteins were highly surface available and inner coat proteins like OxdD seemed to be more surface exposed than would be expected from their documented location. However, our results are consistent with other spore-display studies that have shown many non-crust proteins can result in surface-display of antigens for spore-based vaccines and prior reports that inner proteins like OxdD can load late since spores remain relatively permeable^1,3,6,17–19^. Given than information about SP localization is largely based on analyzing microscopy images of fluorescently tagged proteins, results from our work and that from others highlights the need to re-evaluate the accepted dogma that SPs and SP-cargo fusions are confined to a single coat layer – either natively or during spore-display^8,17,19^. It is possible, and likely, based on our data, that spore coats are heterogeneous, and SPs can localize to multiple layers during assembly. It may also be that spore-display may result in unexpected protein interaction or adsorption and lead to “misincorporation” of the native SP into multiple layers. Given that, our conclusions about localization were internally consistent with the outcomes from the challenge studies. When we challenged 15 of the highest activity SP-GUS fusions to thermal and proteolytic inactivation, we found that most conferred significant resistance to loss in activity, especially compared to the free enzyme. As expected, crust proteins were most susceptible to proteolytic degradation whereas proteins found internally were more stabilizing. Conversely, the crust proteins were some of the most stabilized against thermal inactivation.

We found that SscA was generally poorly surface available when fused to GUS. This, combined with the fact that it is hydrophobic, and its promoter is expressed relatively early, suggests that it may assemble at high copy number within the inner protein layers of spores. Based on these results, our expectations were that surface exposed proteins would be more suitable fusion partners for enzymes that act on insoluble substrates. When we fused DuraPETase to several spore proteins, the highly surface exposed CotY was found to be superior in aiding in the degradation of PET disks than was SscA. However, the SscA-DuraPETase fusion was able to degrade PET (equivalently to another crust protein fusion, CotX), indicating that under some genetic contexts, it may also be surface accessible – again highlighting the putative heterogeneity of spore assembly, especially in genetically engineered contexts.

Our DNA constructs and data on promoter strength and timing, total activity, surface-availability, stability, and combinatorial loading should enable future users to make informed decisions when developing vaccines, biocatalysts, and other products using spore-display. Overall, our extensive survey of spore proteins serves as a key resource for applied scientists and engineers interested in leveraging spores for biotechnological applications.

## AUTHOR CONTRIBUTIONS

Todd C. Chappell, Trevor B. Nicks – Conceptualization; formal analysis; investigation; methodology; data visualization; writing—original & revised draft. Jan-Fang Cheng, Angela Tarver, Jessica A. Lee, Arushi Kalia – Formal analysis; investigation; methodology; data visualization; writing—editing & review. Nikhil U. Nair – Conceptualization; funding acquisition; supervision; writing— original, review & editing.

## ACKNOWLEDGMENTS

We thank all past and present members of the Nair lab for their helpful discussion. We would also like to thank Prof. James A. Van Deventer (Tufts ChBE), Prof. Andrew Camilli (Tufts GSBS), and Prof. Neel Joshi (Northeastern CCB) for their thoughtful feedback on experimental designs. This work was supported by grants DP2HD091798, R21HD105934 (NIH), and CBET-2208390 (NSF) to N.U.N. The work (proposal: https://doi.org/10.46936/10.25585/60001163) conducted by the U.S. Department of Energy Joint Genome Institute (https://ror.org/04xm1d337), a DOE Office of Science User Facility, is supported by the Office of Science of the U.S. Department of Energy operated under Contract No. DE-AC02-05CH11231

## MATERIALS & METHODS

### Bacterial Strains and Culture Conditions

*B. subtilis* 168 (Δ*trpC*) and its derivatives were used for all experiments unless noted otherwise. *E. coli* DH5a or NEB5a were used for general plasmid construction and propagation. General strain propagation was performed in Luria Bertani (LB) at 37 °C and 225 rpm shaking or on LB agar (1.5 %) plates. Chloramphenicol (5 μg mL^−1^) or Kanamycin (10 μg mL^−1^) was added for selection in *B. subtilis* as required and ampicillin (100 μg ml^−1^) for *E. coli*.

### Preparation of Naturally Competent Cells & Transformations

Competent cells were prepared as described previously with slight modifications^20^. A 1 mL aliquot of overnight *B. subtilis* LB culture of was diluted into 15 mL of pre-warmed SM1 medium and incubated at 37 °C 225 rpm until it reached an OD_600_ of 0.5-0.6, (∼4 h). An equal volume of pre-warmed SM2 media was then added and the culture incubated for an additional 90 minutes. Cells were then portioned into 0.5 mL aliquots and used for transformation immediately or stored at -80 °C after adding glycerol to a final concentration of 10 % (v/v). Transformation was performed by adding 100-500 ng of linearized vector DNA to competent cells, incubating for 1 h at 37 °C 225 rpm, and then plating on the appropriate selective media.

### Spore Protein – β-Glucuronidase Fusion Construct Design

N-terminal and C-terminal spore coat protein – B-glucuronidase (SP-GUS) were designed to enable homologous recombination following natural transformation. For N-terminal fusions of GUS to SPs, 2 kb of genomic homology were amplified from the +3-nucleotide position within the coding DNA sequence (CDS) of the SP to insert the GUS fusion at the native locus. The upstream homology arm was amplified with a C-terminal side primer overhang to introduce a flanking AscI cut site between the homology arm and a 6xHis tag sequence. The 6xHis-Tag sequence preceded a 3x(4GS) linker sequence flanked by a NotI site and the GUS sequence. The GUS sequence was flanked by a coding SbfI site and 3x(4GS) linker sequence (no stop codon) that joined its downstream homology arm which includes the SP CDS and additional locus homology. For C-terminal constructs, 2 kb of genomic homology were amplified from the nucleotide preceding the stop codon in each SP. A 3x(4GS) linker was added to the SP CDS which was flanked by an AscI site preceding the GUS CDS. The GUS CDS was flanked by an NotI site, and 3x(4GS) linker to 6xHis tag sequence followed by a SbfI site and the downstream homology arm. For sequences which contained AscI, NotI, or SbfI sites within the CDS or homology arms, other restriction enzyme cut sites were used on a case-by-case basis. All sequences are in **Supplementary Data File 1**. These constructs were then cloned into the pENTR vector backbone. For co-expression analysis of the top five anchors (CgeA, CotG, CotX, CotY, and SscA) the native promoters [defined as the region encompassing 300 bp upstream of the SP CDS start codon] and SP-GUS sequence for each was PCR-amplified from their individual pENTR vector and cloned into pBS1K. These vectors were then linearized and transformed into each of the following *B. subtilis* 168, *B. subtilis* (Δ*sp*), and *B. subtilis* (*sp::sp-gusA*). *B. subtilis* (Δ*sp*) strains were ordered from the BGSC: Δ*cgeA* (BGSC ID: BKE19780), Δ*cotG* (BKE36070), Δ*cotX* (BKE11760), Δ*cotY* (BKE11750), Δ*sscA* (BKE9958).

### Co-Transformation for in-situ Expression of SP-GUS Fusions

Freshly prepared competent cells were portioned into 0.5 mL aliquots, to which 100 ng of linearized selection construct (pBS1K or PBS1C) and 3 μg of linearized pENTR-SP-GUS construct was added and gently mixed by pipetting followed by incubation at 37 °C and 225 rpm for 1 h. Subsequently, 0.5 mL of LB was added to each transformation mixture and incubated for an additional hour. 0.2 mL fractions of the culture were then plated on individual 2×SG sporulation plates containing an antibiotic for the primary marker (pBS1K = 10 μg mL^−1^ Kan, pBS1C = 5 μg mL^−1^ Cm) and X-gluc (5-bromo-4-chloro-3-indolyl-β-D-glucuronic acid, 100 μg mL^−1^). Plates were then incubated at 37 °C for 96 h, during which they were observed for the formation of blue colonies. Blue colonies were colony purified on 2×SG plates and SP-GUS integration was confirmed by colony PCR. 2×SG sporulation plates were composed of 1.6% Difco Nutrient Broth, 0.05 % MgSO_4·_7H_2_O, 0.2 % KCl, 1 mM CaNO_3_, 1 mM MnCl_2_, 0.1 μM FeSO_4_, 0.1 % Glucose, and 1.5 % w/v agar^21^. A X-gluc stock (100 mg mL^−1^ solution in dimethylformamide) was freshly prepared and added to the autoclaved agar to a final concentration of 100 μg mL^−1^.

### Imaging of GUS Activity on Sporulation Plates & Quantification of Promoter Activities

Overnight cultures of SP-GUS construct strains were spotted on 2×SG sporulation plates containing X-gluc. Plates were incubated at 37 °C and images were taken in a lightbox using an iPhone camera at the labeled timepoints. The magnitude of GUS activity was quantified using Fiji by averaging pixel intensity of an ROI defined by colony boundaries after background subtraction using the color deconvolution tool.

### Sporulation & Spore Purification

Sporulation was performed by subculturing LB overnight cultures 1:200 into 2 mL of Difco Sporulation Media (DSM: 0.8 % w/v Tryptone, 0.1 % w/v KCl, 1 mM MgSO_4_, 10 μM MnCl_2_, 1 μM FeSO_4_, 0.5 mM CaCl_2_) and grown for 24 h at 37 °C225 rpm. Spores were harvested after 24 h in DSM through centrifugation at 21,000 ×g for 2 min and washed 3 times with 500 μL water. Following the wash, the top layer of spores was carefully resuspended without disturbing the vegetative cells below and moved to a fresh tube for analysis.

### GUS Activity and Challenge Assays

Spore-displayed GUS activity was quantified spectrophotometrically with the substrate 4-Nitrophenyl β-glucopyranoside (NPG). Reactions were composed of 140 μL reaction buffer (0.1 % v/v β-mercaptoethanol in PBS, pH = 7.4), 30 μL of substrate solution (4 mg mL^−1^ NPG in PBS, pH = 7.4), and 30 μL of spores with a final OD_600_ of 0.1–0.3. Reactions were performed at room temperature with continuous shaking in a 96-well flat bottom plate and measuring the increase in absorbance at 405 nm using a SpectraMax M3 spectrophotometer (Molecular Devices; San Jose, CA). Purified spores resuspended in PBS (pH = 7.4), were incubated at 25, 37, 50, and 60 °C for 15 min in a thermocycler, cooled to room temperature, and tested for activity using the GUS Activity Assayoutlined above. Protease challenge of spores was performed by resuspending spores in PBS containing 0.1-1.0 mg mL^−1^ trypsin, incubating at 37 °C for 1 h and tested for activity using the GUS Activity Assay. GUS with a N-terminal 6xHis tag was cloned into pACYC_Duet1, expressed in *E. coli*, and purified using Ni NTA resin using standard methods to serve as a control for evaluating thermal stability.

### Flow Cytometry

Surface detection of spore-displayed SP-GUS was determined using immunohistochemistry and flow cytometry. Following sporulation, cells were transferred to 50 mL conical flasks and centrifuged at 2,400 ×g for 10 min. The pellet was then transferred to a 1.5 mL Eppendorf and washed 3× with PBS (pH = 7.4) with centrifugation done at 21,000 × g for 2 min. After the final centrifugation, the upper layer of the cell pellet (Figure 3.B) was isolated by gentle pipetting and resuspended in PBSA (2.0 % w/v Bovine Serum Albumin in PBS, pH = 7.4) at an OD_600_ = 1.0 for blocking. Following blocking, 50 μL fractions were pelleted and resuspended in 200 μL of chilled PBSA containing 1 μg mL^−1^ primary antibody (Anti-His tag mouse antibody, Invitrogen, Catalog # MA121315 or Anti-GUS rabbit antibody, Invitrogen, Catalog # A5790) and incubated for 1 h. This was followed by four washes with 200 μL of chilled PBSA and incubation with the secondary antibody in 200 μL of PBSA containing 1 μg mL^−1^ (AlexaFluor 467 goat anti-mouse, Invitrogen, Catalog # A32728 or AlexaFluor 467 goat anti-rabbit, Invitrogen, Catalog # A21245) for 1 h at 4 °C. Samples were then washed four additional times in chilled 200 μL PBSA before being analyzed using an Attune NxT Flow Cytometer (Invitrogen; Carlsbad, CA). 300,000 events were collected for each sample using the lowest possible collection speed, 12.5 μL min^−1^, and five rinse cycles were used between each sample to prevent carryover. Sporulation efficiency of SP-GUS strains was performed by resuspending sporulation cultures with 1x SYBR Safe DNA Stain in PBS and incubating for 30 min in the dark. Following incubation, sporulation cultures were washed 1x with PBS, resuspended in PBS, and immediately analyzed via flow cytometry. Gating for single spore events was based on the FSC-H profile of 1 and 2 μm calibration beads (ThermoFisher, Catalog#: F13838). (**Figure S5**). Gating and analysis were done using FCS Express Software (De Novo Software;Pasadena, California).

### PETase Activity Assays and Imaging

DuraPETase was codon-optimized for *B. subtilis* and ordered as a gene fragment from Twist Biosciences. Standard cloning methods were used to insert DuraPETase into the pBS1K-SP-GUS constructs such that they replace GUS, and these constructs were linearized and integrated into

*B. subtilis*. DuraPETase was also cloned into pACYC_Duet1 with 6xHis tag, expressed in *E. coli*, and purified using Ni-NTA resin using standard methods. To measure the activity of the spore-displayed PETase on a small, soluble molecule we used para-nitrophenol acetate (pNPA, ThermoFisher Catalog #128750050) as a substrate. A pNPA stock solution (100 mg mL^−1^ in DMSO) was diluted to 1.0 mg mL^−1^ in PETase Assay Buffer A (50 mM NaPO_4_, 90 mM NaCl, pH = 7.5). Washed spores were suspended to an OD_600_ of ∼0.1 – 0.25 in PETase Assay Buffer Aand 20 μL of spores were added to 180 μL of PETase Assay Buffer containing 1.0 mg mL^−1^ pNPA pre-aliquoted into a 96-well microplate. Reaction progression was evalulated by tracking formation of the pNP product at 405 nm with a Spetramax M3 spectrophotometer (Molecular Devices, San Jose, CA). Activity of the PETases on insoluble polyethylene terephthalate (PET) discs was evaluated by resuspending purified spores to OD_600_ of ∼1.0 in PETase Assay Buffer B (100 mM KH_2_PO_4_, pH 8.0), adding a single 1.0 cm PET disc, and incubating at 50 °C 225 rpm shaking. After incubating for 48 h, relative PETase activity was measure spectrophotometrically by detecting TPA and MHET product formation at 260 nm. Standard curves of MHET and TPA and purified DuraPETase were used to validate assay conditions (**Figure S7**). PETase treated discs were imaged using a Phenom G2 Purse Tabletop SEM operated at 5 kV. Discs were washed five times with PBS, air-dried overnight, dehydrated with a 5-minute 100 % ethanol bath, sputter coated with a gold-palladium mix prior to imaging.

