## Supplemental Information for "Advancing protein display on bacterial spores through an extensive survey of coat components"

**Table S1.** List of known spore coat proteins, initial documented use in fusion protein display, and their effect on spore function.

| **Coat Protein** | **N or C Fusions** | **Location (If known)** | **Regulate or Effects Incorporation of Others** | **Mutations Effect Sporulation** | **Mutations Effect Function (germination, lysozyme sensitivity)** | **Enzymatic Activity** | **Sources** |
| --- | --- | --- | --- | --- | --- | --- | --- |
| CgeA | C | Crust |  |  |  |  | Imamura, 2010. Imamura, 2014 |
| CmpA |  |  |  |  |  |  | Ebmeier, 2012 |
| CotA |  | Outer Coat |  |  |  | Laccase | Donovan, 1987  Hullo, 2001 |
| CotB | Both | Outer Coat |  |  |  |  | Donovan, 1987  Zilha, 2004. Imamura, 2014 |
| CotC | Both | Outer Coat |  |  |  |  | Donovan, 1987  Isticato, 2004  Imamura, 2014 |
| CotD |  | Outer Coat |  |  |  |  | Imamura, 2010 |
| CotE | C | Outer Coat | Yes |  |  |  | Little, 2001 |
| CotF |  | Inner Coat |  |  |  |  | Cutting, 1991 |
| CotG | Both | Outer Coat |  |  |  |  | Zilha, 2004 |
| CotH |  | Inner Coat | Yes |  | Yes |  | Zilha, 2004 |
| CotI |  | Outer Coat |  |  |  |  | Lai, 2003 |
| CotJA |  |  |  |  |  |  | Seyler, 1997 |
| CotJB |  |  |  |  |  |  | Seyler, 1997 |
| CotJC |  |  |  |  |  |  | Seyler, 1997 |
| CotM |  | Outer Coat |  |  |  |  | Henriques, 1997 |
| CotN |  |  | Yes |  |  |  | Stover, 1999 |
| CotO |  |  | Yes |  | Yes |  | McPherson, 2005 |
| CotQ |  |  |  |  |  | Oxidoreductase | Lai, 2003 |
| CotS |  | Outer Coat |  |  |  |  | Takamatsu, 1998 |
| CotT |  | Inner Coat |  |  | Yes |  | Imamura, 2010 |
| CotU |  | Outer Coat |  |  |  |  | Lai, 2003 |
| CotV | C | Crust |  |  |  |  | Krajcikova, 2009 |
| CotW |  | Crust |  |  |  |  | Krajcikova, 2009 |
| CotX | C | Crust |  |  |  |  | Zhang, 1993 |
| CotY | C | Crust |  |  |  |  | Zhang, 1993 |
| CotZ | C | Crust |  |  |  |  | Zhang, 1993 |
| CwIJ |  | Inner Coat |  |  |  | Cell Wall Hydrolase | Ishikawa, 1998 |
| GerQ |  | Inner Coat |  |  |  |  | Ragkousi, 2003 |
| LipC |  | Basement |  |  |  | Phospholipase | Masayama, 2007 |
| OxdD | C | Inner Coat |  |  |  | Decarboxylase | Potot, 2010 |
| SafA |  | Inner Coat |  |  | Yes |  | Ozin, 2001 |
| SodA |  |  |  |  |  | Superoxidase Dismutase | Henriques, 1998 |
| SpoIVA |  | Basement | Yes |  |  |  | Roels, 1992 |
| SpoVID |  | Basement | Yes |  |  |  | Beall, 1993 |
| SpoVIF |  |  |  |  | Yes |  | Kuwana, 2003 |
| SscA |  |  |  |  | Yes |  | Kudamo, 2011 |
| Tgl |  | Inner and Outer Coat |  |  |  | Transglutaminase | Kobayashi, 1998 |
| SleL |  | Inner Coat |  |  |  | N-acetylglucosaminidase | Ustock, 2015 |
| YabG |  | Outer Membrane |  |  |  | Cysteine Protease | Takamatsu, 2000 |
| YabP |  | Outer Membrane |  |  |  |  | Ooij, 2004 |
| YbaN |  |  |  |  |  | Polysaccharide Deacetylase | Silvaggi, 2004 |
| YeeK |  | Inner Coat |  |  |  |  | Takamatsu, 2009 |
| YhaX |  | Basement |  |  |  |  | Eichenberger, 2003 |
| YhbB |  |  |  |  |  | Amidase | Eichenberger, 2003 |
| YheD |  | Basement |  |  |  |  | Ooij, 2004 |
| YhjR |  | Inner Coat |  |  |  |  | Kuwana, 2002 |
| YknT |  | Outer Coat |  |  |  |  | Eichenberger, 2004 |
| YmaG |  | Inner Coat |  |  |  |  | Kuwana, 2002 |
| YppG |  | Basement |  |  |  |  | Eichenberger, 2004 |
| YsnD |  | Inner Coat |  |  |  |  | Kuwana, 2002 |
| YtrH |  |  |  | Yes |  |  | Eichenberger, 2003 |
| Ytrl |  | Membrane |  | Yes |  |  | Eichenberger, 2003 |
| YtxO |  | Outer Coat |  |  |  |  | Imamura, 2010 |
| YutH |  | Inner Coat |  |  |  |  | Ooij, 2004 |
| YuzC |  | Inner Coat |  |  |  |  | Eichenberger, 2003 |
| YxeE |  | Inner Coat |  |  |  |  | Steil, 2005 |

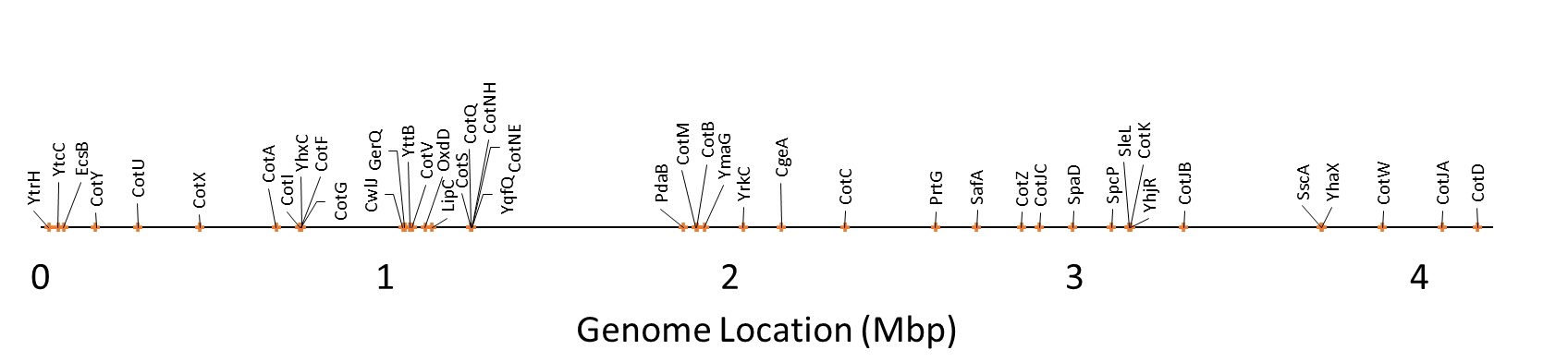

**Figure S1.** Genomic locations of all 44 spore coat proteins under investigation.

**Table S2.** List of constructs that were successfully validated at different characterization steps.

| **Starting Constructs (52)** | **PCR-Confirmed (34)** | **Functionally Spore-Displayed (32)** |
| --- | --- | --- |
| CgeA-C-GusA-His | CgeA-C-GusA-His | CgeA-C-GusA-His |
| CgeA-N-GusA-His | CgeA-N-GusA-His | CgeA-N-GusA-His |
| CotA-C-GusA-His | CotA-C-GusA-His | CotA-C-GusA-His |
| CotA-N-GusA-His |  |  |
| CotB-C-GusA-His | CotB-C-GusA-His | CotB-C-GusA-His |
| CotB-N-GusA-His |  |  |
| CotC-C-GusA-His | CotC-C-GusA-His | CotC-C-GusA-His |
| CotC-N-GusA-His | CotC-N-GusA-His | CotC-N-GusA-His |
| CotD-C-GusA-His |  |  |
| CotG-C-GusA-His | CotG-C-GusA-His | CotG-C-GusA-His |
| CotG-N-GusA-His | CotG-N-GusA-His | CotG-N-GusA-His |
| CotI-C-GusA-His | CotI-C-GusA-His | CotI-C-GusA-His |
| CotI-N-GusA-His | CotI-N-GusA-His | CotI-N-GusA-His |
| CotJA-C-GusA-His | CotJA-C-GusA-His | CotJA-C-GusA-His |
| CotM-C-GusA-His | CotM-C-GusA-His | CotM-C-GusA-His |
| CotQ-N-GusA-His |  |  |
| CotS-C-GusA-His | CotS-C-GusA-His | CotS-C-GusA-His |
| CotS-N-GusA-His |  |  |
| CotU-C-GusA-His | CotU-C-GusA-His | CotU-C-GusA-His |
| CotU-N-GusA-His |  |  |
| CotV-C-GusA-His | CotV-C-GusA-His | CotV-C-GusA-His |
| CotV-N-GusA-His |  |  |
| CotW-C-GusA-His | CotW-C-GusA-His | CotW-C-GusA-His |
| CotW-N-GusA-His | CotW-N-GusA-His | CotW-N-GusA-His |
| CotX-C-GusA-His | CotX-C-GusA-His | CotX-C-GusA-His |
| CotY-C-GusA-His | CotY-C-GusA-His | CotY-C-GusA-His |
| CotY-N-GusA-His | CotY-N-GusA-His | CotY-N-GusA-His |
| EcsB-C-GusA-His | EcsB-C-GusA-His | EcsB-C-GusA-His |
| LipC-N-GusA-His |  |  |
| OxdD-C-GusA-His | OxdD-C-GusA-His | OxdD-C-GusA-His |
| OxdD-N-GusA-His | OxdD-N-GusA-His | OxdD-N-GusA-His |
| PdaB-C-GusA-His | PdaB-C-GusA-His |  |
| PdaB-N-GusA-His | PdaB-N-GusA-His |  |
| PrtG-C-GusA-His | PrtG-C-GusA-His | PrtG-C-GusA-His |
| PrtG-N-GusA-His |  |  |
| SafA-N-GusA-His |  |  |
| SleL-C-GusA-His |  |  |
| SpaD-C-GusA-His | SpaD-C-GusA-His | SpaD-C-GusA-His |
| SpaD-N-GusA-His |  |  |
| SscA-C-GusA-His | SscA-C-GusA-His | SscA-C-GusA-His |
| SscA-N-GusA-His |  |  |
| YhaX-C-GusA-His | YhaX-C-GusA-His | YhaX-C-GusA-His |
| YhaX-N-GusA-His |  |  |
| YhjR-N-GusA-His | YhjR-N-GusA-His | YhjR-N-GusA-His |
| YhxC-N-GusA-His |  |  |
| YmaG-N-GusA-His |  |  |
| YqfQ-C-GusA-His | YqfQ-C-GusA-His | YqfQ-C-GusA-His |
| YrkC-C-GusA-His | YrkC-C-GusA-His | YrkC-C-GusA-His |
| YrkC-N-GusA-His |  |  |
| YtcC-C-GusA-His | YtcC-C-GusA-His | YtcC-C-GusA-His |
| YtcC-N-GusA-His |  |  |
| YtrH-N-GusA-His | YtrH-N-GusA-His | YtrH-N-GusA-His |

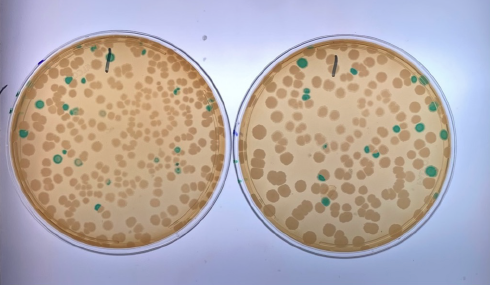

**Figure S2. Representative co-transformation sporulation plates.** Colonies from the co-transformation of linearized pBS1K + pENTR-CgeA-C-GUS and plating on 2xSG media + X-gluc. Blue colonies are GUS^+^ and underwent colony purification and colony PCR to confirm integration.

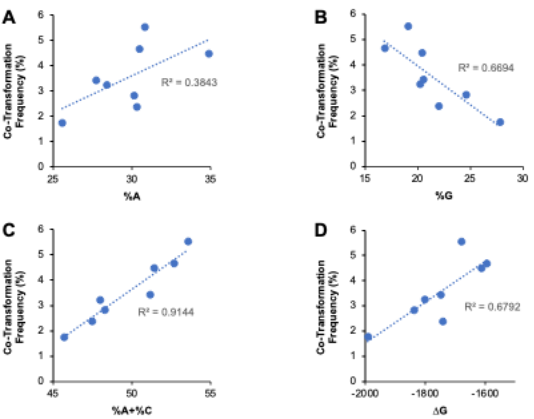

**Figure S3. Co-transformation correlational relationships.** Scatter plot and linear regression analysis of co-transformation frequency and **A)** adenine content (%A), **B)** guanine content (%G), **C)** adenine + cytosine content (%A + %C), and **D)** free energy of folding of the GUS insert and 2 kb flanking arms. R^2^ values are provided for trend lines to indicate degree of fit.

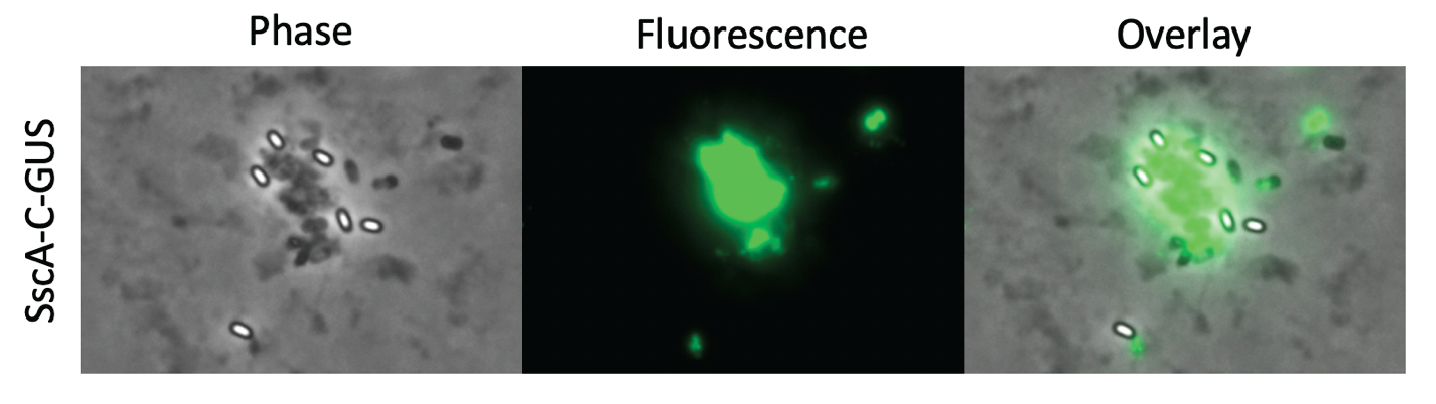

**Figure S4.** Debris from lysozyme treated cells form particles that are hard to distinguish from spores. When using the common lysozyme treatment for cell-lysis at the end of sporulation, we found it difficult to separate spores from debris for immunostaining and flow cytometry. Here is an example of lysed cells and spores carrying SscA-C-GUS. Phase-bright spores (white ovals with black outline) are not individually stained following immunostaining with the anti-HIS antibody. Rather, fluorescence is localized to clumps of cell-debris. To enable accurate immunostaining, we began separating spores from vegetative cells through differential centrifugation.

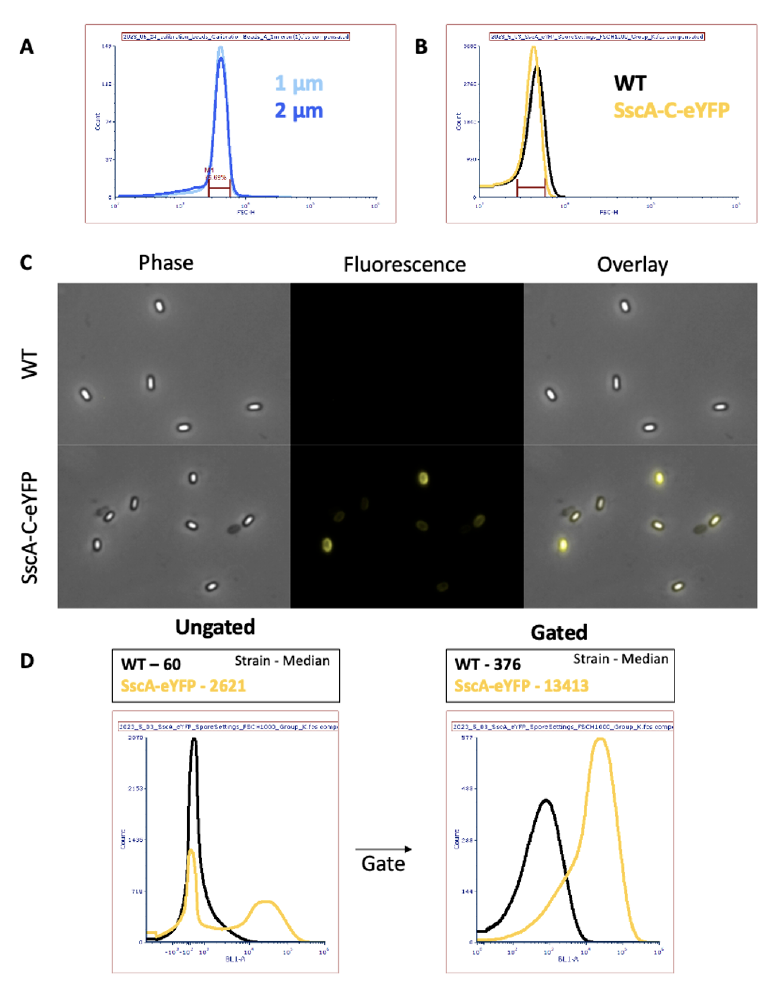

**Figure S6.** Size calibration for spore-based flow cytometry events. **A)** FSC-H histograms of the 1 μm and 2 μm beads essentially overlapped. We created a gate (maroon) based on the peak of these distributions that encompassed ~75% of all events between the two bead sizes. **B)** FSC-H profiles of WT and SscA-C-eYFP spores purified through the method shown in Figure 4. The same gate overlapped with most of the events from both WT and SscA-C-eYFP spore populations. **C)** Fluorescence microscopy images of the purified WT and SscA-C-eYFP spores show no clumping and loading of SscA-C-eYFP to the spore. **D)** Applying the FSC-H gate to the WT and SscA-C-eYFP population yields two distinct fluorescence (BL1-H) peaks: a WT peak with a mean value of 376 and a SscA-eYFP peak with a mean value of 13,413.

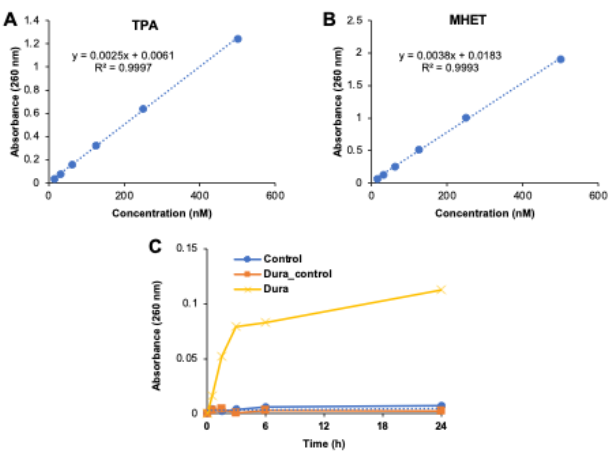

**Figure S7.** **PETase assay validation. Standard curves for PETase products. A)** TPA and **B)** MHET. Both products absorb at 260 nm (A_260_). **C)** Control experiment showing that without PETase and with PET disc (control), or with PETase and without PET disc (Dura_Control) so change in A_260_ is observed, while with both PETase and PET disc (Dura) TPA and MHET product formation are detectable.
